# *Papiliotrema maritimi* f.a. sp. nov., a new tremellaceous yeast species associated to macrophytes in a Marshland of South Brazil

**DOI:** 10.1101/2020.06.15.153189

**Authors:** Mauricio Ramírez-Castrillón, Fernanda Fraga Gomes, Andrea Formoso de Souza, Belize Rodrigues Leite, Danielle Machado Pagani, Patricia Valente

## Abstract

One new yeast species, *Papiliotrema maritimi* sp. nov., is being proposed to be suitable into the Rhynchogastremataceae family, belonging to the Tremellales clade. This new species is related to six others from the *Papiliotrema* genus: *P. taeanensis, P. siamense, P. perniciosus, P. nemorosus, P. bandonii, P. japonica* and *P. fuscus*. The novel species is proposed based on the phylogenetic species concept through analysis of the D1/D2 region, part of the large subunit (LSU) rRNA gene and the internal transcribed spacer (ITS) region. A total of 11 strains of *Papiliotrema maritimi* sp. nov. were obtained from macrophytes leaves collected in south Brazil. *Papiliotrema maritimi* sp. nov. differs by 12, 15, 25, 25, 25 and 29 substitutions in the D1/D2 domain from the related *species P. fuscus, P. japonica, P. siamense, P. nemorosus, P. bandonii, and P. perniciosus*, respectively. Concerning the ITS region, there are 11 substitutions and 52 or more substitutions when compared to *P. teanensis* and its closest relatives. The type strain of *Papiliotrema maritimi* sp. nov. is UFMG-CM-Y6048. The MycoBank number for *Papiliotrema maritimi* sp. nov. is MB 835603.

## Introduction

Basidiomycetous yeasts were recently re-classified inside the Tremellomycetous group according to Liu et al. (2015). Within the family Rhynchogastremataceae, the genera *Rhynchogastrema* and *Papiliotrema* have been proposed. Based on our phylogenetic analyses, we propose a new species of the genus *Papiliotrema* based on eleven yeast strains. These strains were isolated from leaves of three plant species associated to a marshland in the city of Rio Grande, Rio Grande do Sul, Brazil, and represented one undescribed species with variable physiological profiles and almost identical sequences of the ITS and LSU regions among strains (Souza et al. 2014). Analysis of the ITS and LSU sequences showed that this new species differed significantly from other species inside the *Papiliotrema* clade, such as *Papiliotrema aureus, P. flavescens, P. terrestris, P. baii, P. ruineniae, P. wisconsinensis, P. plantarum* and *P. pichitensis*. In this sense, we propose *Papiliotrema maritimi* sp. nov. to be characterized by these yeast strains.

Five samplings were made between June 2012 and January 2013, comprising fresh and decomposing leaf samples. The decomposing macrophytes were stored in litter bags and placed back in the salt marsh, being removed after 7, 14, 40 and 100 days of decomposition to isolate of the yeasts. Samplings, both from litter bags and fresh macrophytes, took place on June 25, July 2, July 30 and September 29 of 2012. All the samples were placed in sterile polyethylene bags and immediately processed at Universidade Federal do Rio Grande do Sul, Porto Alegre, Brazil, washed with sterile distilled water, 3 g of leaves were added to 30 mL of Tween 20 0.5% (v/v) by 200 rpm and 30 min. Subsequently, serial dilutions of the solution were inoculated on YM culture medium (1 % glucose, 0.3 % malt extract, 0.3 % yeast extract, 0.5 % peptone, 2 % agar, pH 4), containing chloramphenicol 200 mg/L, and incubated at 20–25 °C for up to 7 days.

All yeast isolates were purified via the method of repeated streaking onto YM or GPY (2 % glucose, 0.5 % yeast extract, 1 % peptone, 2 % agar) agar plates and preserved in sterile mineral oil at 4°C and GPY medium with sterile glycerol 30% (v/v) at −20°C. for later identification. The yeasts were characterized by standard methods (Kurtzman et al., 2011). Carbon and nitrogen assimilation assays were carried out in solid plates. Details concerning the yeast isolates are reported in Table 1. Strains BEL43, BEL45, BEL46, BEL80, BEL84, BEL87, BEL89, BEL90 and BEL108 were obtained from fresh leaves of *Bolboschoenus maritimus.* Strains DEC01 and DEC07 were isolated from decomposing leaves (7^th^ day) of *Spartina alterniflora* and *S. densiflora*, respectively. All three are macrophyte plant species that normally grow in “Ilha da Polvora” (32°01’20.8”S 52°06’18.0”W), “Lagoa dos Patos”, city of Rio Grande, in the Rio Grande do Sul state, South Brazil.

**Table 1.**
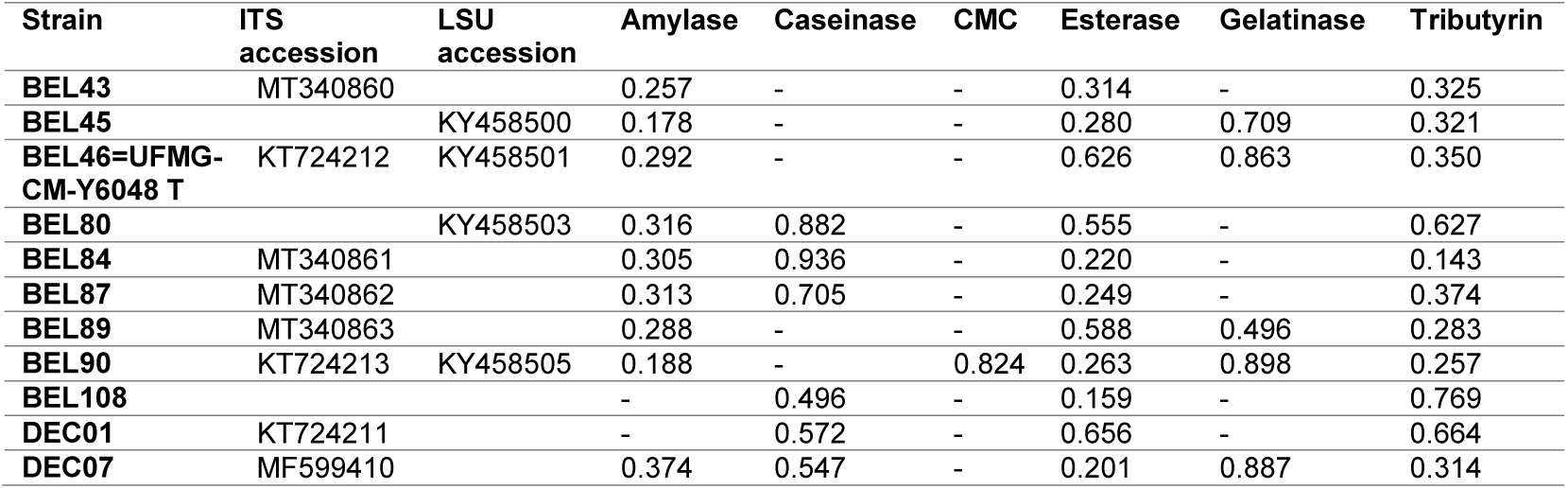
Genbank accession codes and enzymatic activity (represented by Pz values for each test) for each strain of *Papiliotrema maritimi* f.a. sp. nov. – indicates negative result.

| Strain | ITS accession | LSU accession | Amylase | Caseinase | CMC | Esterase | Gelatinase | Tributyrin |
| --- | --- | --- | --- | --- | --- | --- | --- | --- |
| BEL43 | MT340860 |  | 0.257 | - | - | 0.314 | - | 0.325 |
| BEL45 |  | KY458500 | 0.178 | - | - | 0.280 | 0.709 | 0.321 |
| BEL46=UFMG-CM-Y6048 T | KT724212 | KY458501 | 0.292 | - | - | 0.626 | 0.863 | 0.350 |
| BEL80 |  | KY458503 | 0.316 | 0.882 | - | 0.555 | - | 0.627 |
| BEL84 | MT340861 |  | 0.305 | 0.936 | - | 0.220 | - | 0.143 |
| BEL87 | MT340862 |  | 0.313 | 0.705 | - | 0.249 | - | 0.374 |
| BEL89 | MT340863 |  | 0.288 | - | - | 0.588 | 0.496 | 0.283 |
| BEL90 | KT724213 | KY458505 | 0.188 | - | 0.824 | 0.263 | 0.898 | 0.257 |
| BEL108 |  |  | - | 0.496 | - | 0.159 | - | 0.769 |
| DEC01 | KT724211 |  | - | 0.572 | - | 0.656 | - | 0.664 |
| DEC07 | MF599410 |  | 0.374 | 0.547 | - | 0.201 | 0.887 | 0.314 |

The divergent D1/D2 domains of the LSU rRNA gene were amplified with the NL1 and NL4 primers and the ITS-5.8S region with ITS5 and ITS4 primers. DNA of the strains were amplified as described by Ramirez-Castrillon et al. (2017), purified with ExoSapIt and sequenced at “Unidade de Análises Moleculares e de Proteínas (UAMP), Hospital de Clínicas de Porto Alegre”, Brazil, using standard protocols. The obtained sequences were edited, and the consensus sequence were compared with the GenBank database using the BLAST algorithm (Zhang et al., 2000). To estimate phylogenetic relationships, Maximum Likelihood trees of concatenated sequences of ITS and LSU regions were generated with General Time Reversible model (Nei and Kumar 2000) using MEGA X software (Kumar et al., 2018). The analysis involved 39 nucleotide sequences and a total of 1168 positions were performed in the final dataset. The robustness of trees was calculated with 1000 bootstrap pseudo replicates. All analyzed sequences were taken from GenBank database and accession numbers are shown in Figure 1.

**Figure 1.**
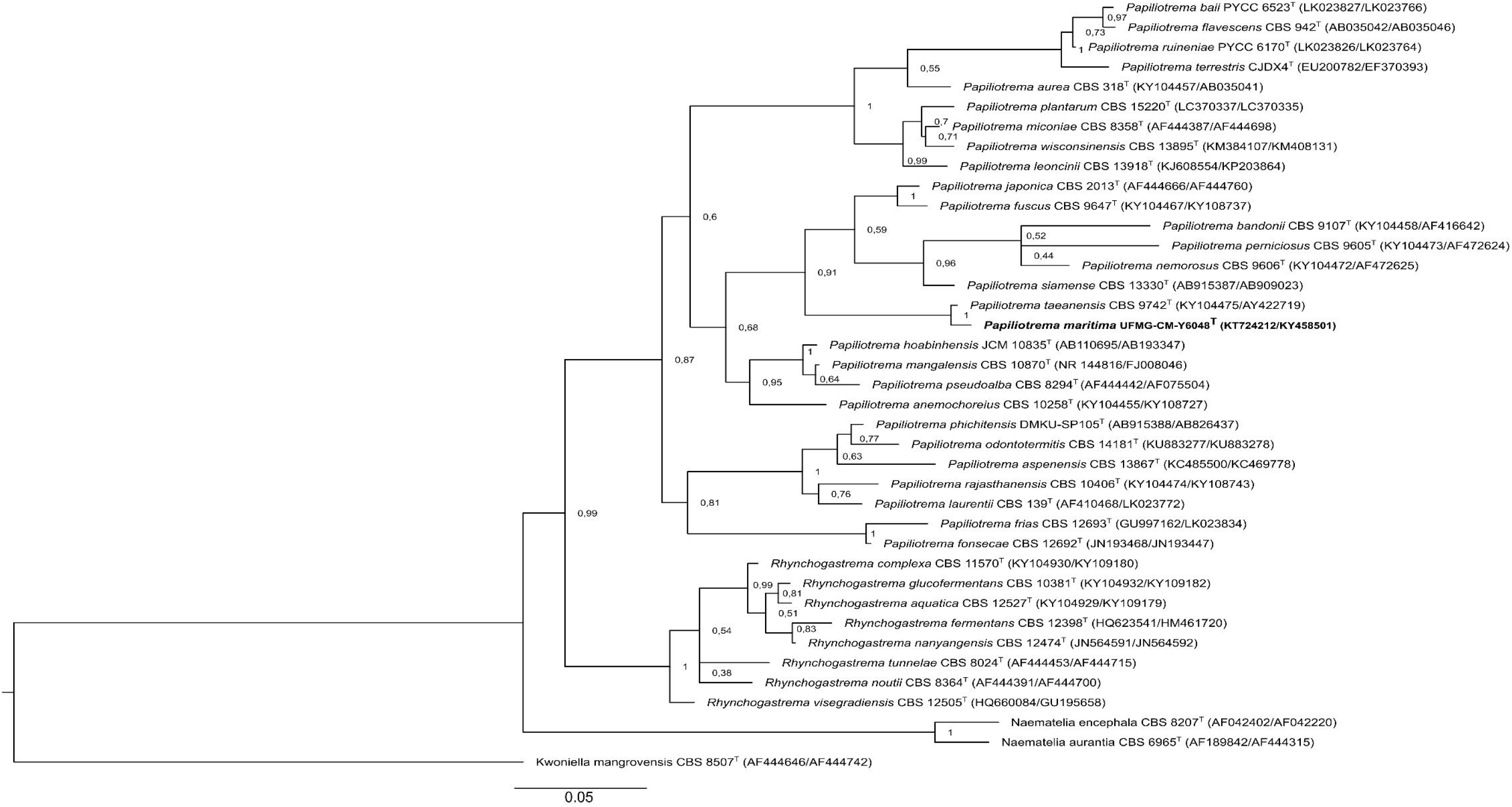
Phylogenetic placement of *Papiliotrema maritimi* f.a. sp. nov. within the Rhynchogastremataceae family as obtained by Maximum Likelihood method (Nei and Kumar 2000) analysis of the concatenated D1/D2 (LSU) and ITS regions. The best model was General Time Reversible (+G +I). Bootstrap values are shown (1000 pseudo replicates). Bar, 0.05 substitutions per nucleotide position. The analysis involved 39 nucleotide sequences with *Kwoniella mangrovensis* CBS8507^T^ as outgroup. There were a total of 1168 positions in the final dataset. Evolutionary analyses were conducted in MEGA X (Kumar et al. 2018).

According to Liu et al. (2015), *Papiliotrema bandonii* is the type species of the genus, and together with *P. fuscus* and *P. plantarum*, have sexual states, which form clavate basidia with transverse septae. However, the sexual state of *P. bandonii* has only been observed *in natura* (Kurtzman et al. 2011). Kurtzman (1973) reported sexual activity in strains of *P. laurentii*. In addition, Yurkov et al. (2015) detected genetic recombination and the presence of sexual reproduction-related genes (MAT genes) in *P. flavescens* and *P. terrestris*, although they failed to induce mating. The authors suggested that their failure could be due to unknown specific media components or environmental conditions to induce sexual reproduction. According to our results, sexual reproduction was not observed in the new species.

### Proposal of a new anamorph species

Delineation of the novel species was based primarily on the analysis of the concatenated sequences of the genes encoding the ITS region and the D1/D2 region of the LSU rRNA gene. A phylogenetic analysis demonstrated that strain UFMG-CM-Y6048 formed a subclade with the other *Papiliotrema* species in the bandonii clade, but the strain UFMG-CM-Y6048 was placed in a clear different position from the other *Papiliotrema* species (Fig. 1), which confirmed its recognition as a novel species.

Sequence comparisons of the ITS region and the D1/D2 domains of the LSU rRNA gene indicated that the 11 Brazilian strains belong to a novel anamorphic yeast species in the *Papiliotrema* genus, and their closest relatives are *P. taeanensis, P. siamense, P. perniciosus, P. nemorosus, P. bandonii, P. japonica* and *P. fuscus* (Fig. 1). All 11 strains were obtained from leaves, collected in a marshland of “Lagoa dos Patos”, south Brazil. Even though the D1/D2 domain (LSU) of the proposed new species is identical to *P. taeanensis*, the ITS region differed by 11 substitutions. The proposed new species differed 25 substitutions on the LSU region and 52 substitutions on the ITS region from *P. siamense*; 25 and 57 from *P. nemorosus*, 29 and 59 from *P. perniciosus*, 25 and 64 substitutions from *P. bandonii*, 15 and 55 substitutions from *P. japonica*, and 12 and 51 from *P. fuscus*, respectively. Pairwise comparisons of sequences demonstrated that all strains of *Papiliotrema maritimi* f.a. sp. nov. had more than 99 % identity in the D1/D2 region (0–1 substitution and 0–1 indel) and identical sequence on the ITS region.

Although formations of basidiospores were not observed, in conformity with the International Code of Nomenclature for Algae, Fungi and Plants, anamorphic and teleomorphic species can be assigned to the same genus (McNeill et al. 2012); the species is therefore assigned to the genus *Papiliotrema*. The designation *forma asexualis* (f.a.) was added to indicate the lack of a sexual cycle in the species assigned to the genus (Lachance 2012). Therefore, we propose the novel species named *Papiliotrema maritimi* f.a. sp. nov. to accommodate these isolates.

Phenotypically, *Papiliotrema maritimi* sp. nov. is very similar to other related species in the bandonii clade, however, it can be distinguished from these species based on nitrate assimilation (delayed), since other species, such as *P. taeanensis, P. fuscus and P. japonica*, are negative (Sampaio et al. 2004; Shin et al. 2005). Many enzymes of biotechnological interest were qualitatively tested and showed variability inside the species (Table 1) according to the Pz value (equation 1), where the produced halo showed promising values for caseinase and gelatinase activities. The strains were tested for lipid accumulation and the results showed no accumulation after five days of growth in A medium (Ramirez-Castrillon et al. 2017). Finally, when the strains were characterized for carbon or nitrogen assimilation, we noted variability in the colony pigmentation, being clearer or darker, depending on the carbon and nitrogen source (Figure 2). For this, we photographed each colony after seven days of growth for each carbon or nitrogen assimilation test. After, we extracted the RGB color code and constructed a heatmap graph using a correlation matrix.

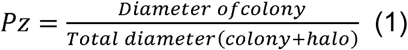

**Figure 2.**
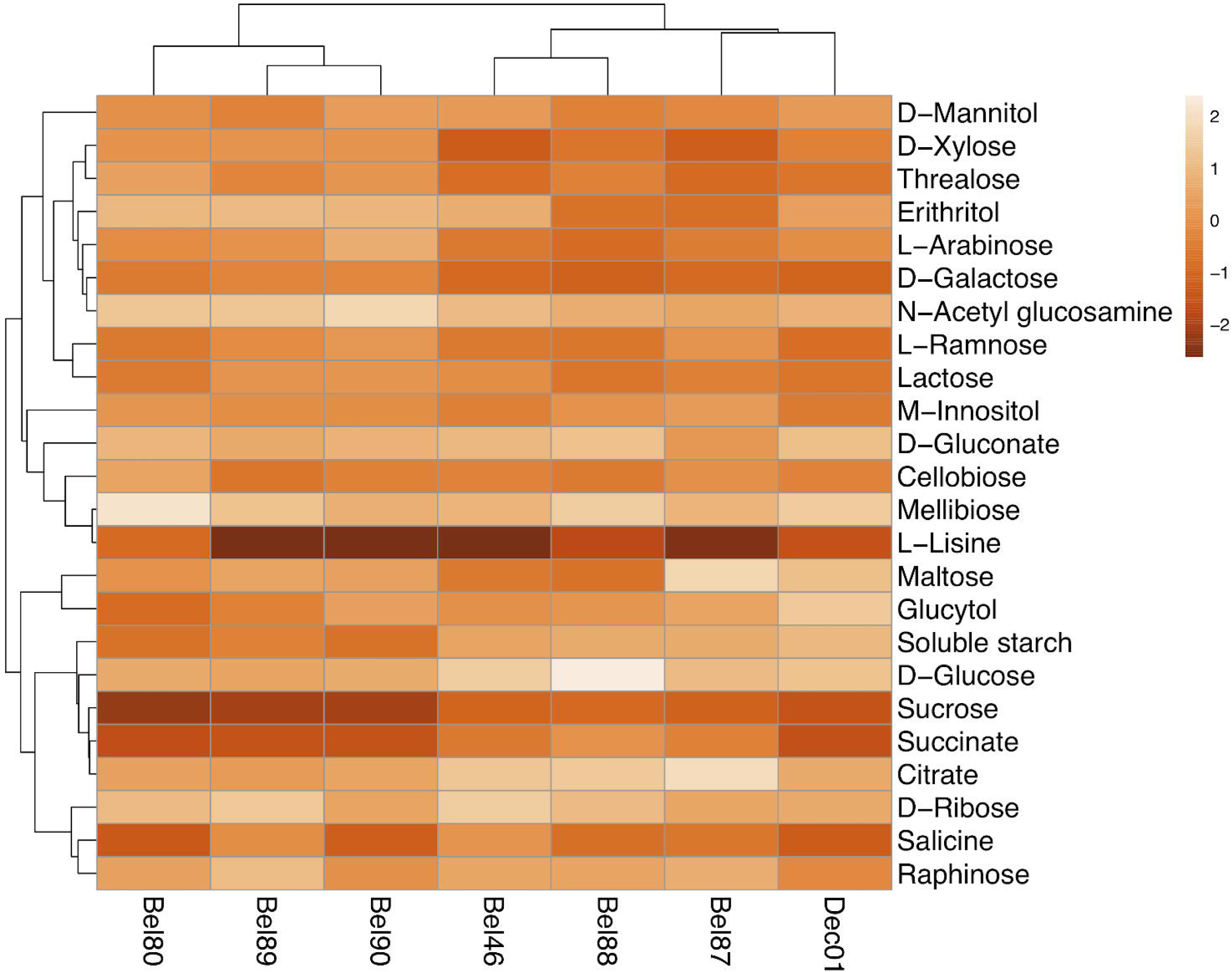
Heatmap showing variability of pigmentation of the colony, depending on the carbon or nitrogen assimilation test. Each colony was photographed and determined their RGB color code. After that, all the values were added to the Clustvis software (Metsalu et al. 2015), where the heatmap was constructed, according to the variances of each value, being values darker or lighter very different to the total average value (orange). Dendrograms were also constructed to group very similar strains or test with similar results using Pearson’s correlation and average linkage method.

### Description of *Papiliotrema maritimi* sp. nov. Ramirez-Castrillon, Formoso, Gomes, Leite, Pagani & Valente

*Papiliotrema maritimi* (ma.ri.ti.mi gen. fem. n. *maritimi* refers to the botanical species name of *Bolboschoenus maritimus*, isolated from the leaf of *Bolboschoenus maritimus*).

In GPY broth after 5 days at 25 °C, the yeast cells are oval to ellipsoid and occur singly, in pairs or multipolar budding (Fig. 2). The colonies are smooth, mucous to butyrous, glistening, cream to pink color, and with an entire margin. Sexual reproduction was not observed after growth in Acetate agar. Ballistoconidia production is absent. Glucose fermentation ability is negative. The following carbon compounds are assimilated: D-glucose, galactose, D-ribose (delayed or d), D-xylose, arabinose, L-rhamnose, sucrose, maltose (d), trehalose, soluble starch, glycerol (d), erythritol, cellobiose, salicin, melibiose, lactose, raffinose, D-mannitol, M-inositol, D-gluconate, succinate (d), citrate (variable), N-acetyl-glucosamine, sorbitol. No growth was observed on methanol, acetone, isopropanol, tween 80, and ethanol. The following nitrogen compounds were assimilated: sodium nitrate (d) and L-lysine; no growth was observed on creatine and sodium nitrite. Growth at 30 °C is positive. Growth was not observed on GPY with 10% or 15% sodium chloride, 0.01% or 0.1% cycloheximide. When grown in different carbon or nitrogen sources, some strains showed variability in pigmentation, as showed in Figure 3. The type strain was isolated from the leaf of *B. maritimus* in a marshland located in “Lagoa dos Patos”, Rio Grande, RS, Brazil. The type strain was deposited in the collection of Microorganisms and Cells at Universidade Federal de Minas Gerais, Belo Horizonte, MG, Brazil, as strain UFMG-CM-Y6048.

**Figure 3.**
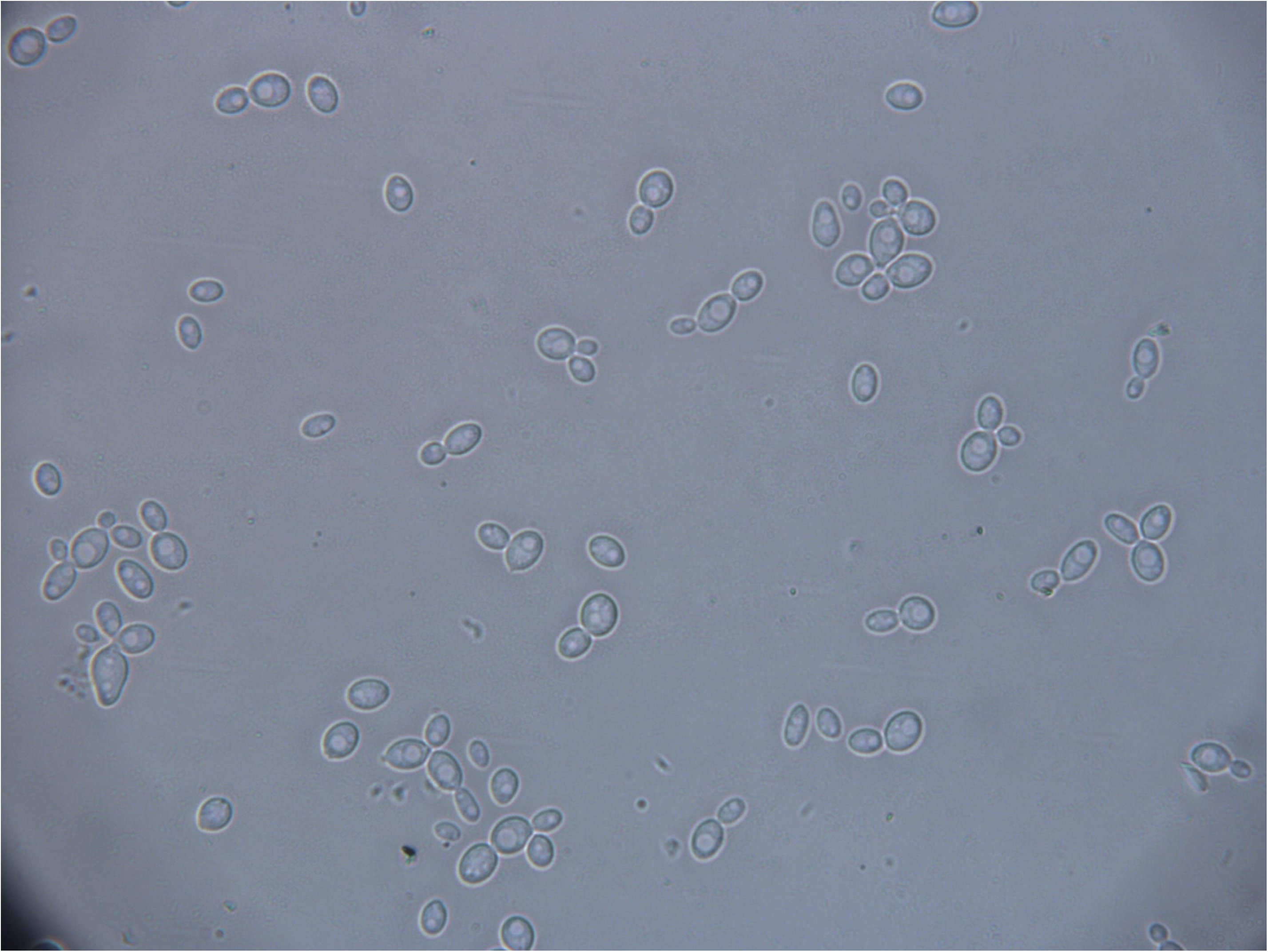
Cell morphology of *Papiliotrema maritimi* UFMG-CM-Y6048 after 48h of growth in GPY broth. 1000X magnification.

The MycoBank number is MB 835603.

## Acknowledgements

This work was supported by *Conselho Nacional de Desenvolvimento Científico e Tecnológico* (CNPq, Brazil), *Coordenação de Aperfeiçoamento de Pessoal de Nível Superior* (CAPES, Brazil), *Fundação de Amparo à pesquisa do Estado do RS* (FAPERGS, Brazil), Universidad Santiago de Cali (Colombia) [Grant number 934-621118-8], and *Ministerio de Ciencia, Tecnologia e Innovación* MINCIENCIAS (Colombia) [Grant numbers 512, 784].

